# Assessing Computational Models for Pharmacogenomic Variant Interpretation

**DOI:** 10.64898/2026.08.03.742561

**Authors:** Fabrizio Pucci, Pauline Hermans, Matsvei Tsishyn, Jessica Cusato, Marianne Rooman

## Abstract

Accurately predicting the effects of pharmacogenomic variants is essential for the development of personalized therapeutic strategies, as genetic variability can influence drug response differently across patients. Here, we assessed several computational approaches using a dataset of pharmacogenomic variants with either clinical annotations or functional characterization by deep mutational scanning, compiled from the literature, with an additional focus on CYP2C9, a clinically relevant drug-metabolizing enzyme. Our results show that, despite recent methodological advances, substantial room for improvement remains. In particular, current methods struggle to distinguish gain-of-function variants associated with increased drug clearance and fast-metabolizer phenotypes from neutral variants, whereas loss-of-function variants that reduce drug clearance are predicted more accurately. The integration of structural and evolutionary information appears to be a key strategy for improving performance, with the coevolution-based StructureDCA method achieving the highest accuracy compared with classical genetic variant-effect predictors and recent deep learning approaches, including the pathogenic-variant predictor AlphaMissense and general protein language model–based methods. Finally, our results indicate that computational models can complement in vitro experiments in clinical variant interpretation, as StructureDCA predictions showed better agreement with clinically annotated phenotypes than large-scale deep mutational scanning data in several cases.

## 1. Introduction

Inter-individual genetic differences are a key determinant of variability in both drug efficacy and toxicity, contributing to heterogeneous treatment outcomes and adverse drug reactions.^1,2^ Developing reliable methods to infer drug response directly from a patient’s genomic profile would represent a major step toward precision medicine. Such approaches could support more individualized therapeutic decisions, improve treatment effectiveness and safety, as well as reduce healthcare costs.^3^

Genomic variants of pharmacogenomic relevance have already been incorporated into clinical guidelines. A classical example is warfarin, for which polymorphisms in n the drug-metabolizing enzyme CYP2C9, the pharmacological target VKORC1, and, to a lesser extent, the vitamin K–metabolizing enzyme CYP4F2 contribute to interindividual variability in dose requirements. These genetic variants are routinely considered in pharmacogenetic dosing algorithms and are incorporated into clinical guidelines.^4^

Another example of pharmacogenomics in oncology is fluoropyrimidine therapy, where genetic variants in the diphyrimidine dehydrogenase gene (DPYD) are used to guide treatment. Patients carrying decreased-function DPYD alleles have a markedly increased risk of severe or life-threatening toxicity following treatment with fluorouracil or capecitabine. Consequently, the guidelines recommend genotype-guided dose reductions or the use of alternative therapies for these patients.^5^

However, despite significant advances in genomics, linking genetic variation to drug response is still a major challenge in clinical pharmacology.^6^ Indeed, genetic variants can influence drug response through a wide range of pharmacokinetic and pharmacodynamic mechanisms.^7^ From a pharmacokinetic perspective, genetic variants can affect the four major ADME processes : drug absorption (A), distribution (D), metabolism (M), and elimination (E), by altering the function or expression of drug transporters, metabolizing enzymes, and other proteins involved in these processes. From a pharmacodynamic perspective, genetic variants can affect drug target affinity and expression, intracellular signaling pathways, immune recognition, and other biological processes that determine the cellular response to a therapeutic agent. Together, the variety of these mechanisms highlights the complexity of the inter-individual variability in drug efficacy and toxicity, as well as the challenges of understanding and predicting drug response from genomic information alone. To complement the time-consuming and costly experiments in pharmagenomic variant characterization, computational methods have been applied to get insight into the impact of the selected variants.

Over the past decades, substantial efforts have been devoted to developing computational models for predicting the functional effects of genetic variants, ranging from classical methods such as PolyPhen-2,^8^ PROVEAN^9^ and SIFT^10^ to more recent deep learning–based approaches. The scientific community has also invested considerable effort in evaluating these methods through systematic benchmarks and blind challenges. In this context, the Critical Assessment of Genome Interpretation (CAGI) has played a major role in advancing the field by providing rigorous and independent assessments of variant-effect predictors.^11,12^

In this study, we evaluated state-of-the-art variant-effect predictors for assessing the functional consequences of pharmacogenomic variants, including deep learning–based approaches such as AlphaMissense,^13^ several protein large language models (pLLMs) such as ESM-2,^14^ and our recently published in-house tools RSALOR^15^ and StructureDCA,^16^ compared to established classical variant-effect predictors. All methods were assessed on curated pharmacogenomic variant datasets derived from clinical annotations or high-throughput in vitro experimental assays.

## 2. Methods

### 2.1. Functional data collection

To benchmark state-of-the-art methods for predicting the functional effects of genomic variants, we curated three pharmacogenomic variant datasets, on which none of the evaluated predictors have been trained.

The first dataset, *D*_CL_, was obtained from^17^ and comprises 337 variants across 43 genes. For each variant, the pharmacogenomic phenotype was represented by the mean in vitro functional activity, expressed as the intrinsic clearance (CLint) of the mutant protein relative to the corresponding wild-type protein (%), and measured using one or, in some cases, two substrates. The analyzed genes primarily belong to the ADME functional family and include genes encoding phase I drug-metabolizing enzymes (e.g., CYP1A2, CYP1B1, CYP2A6, CYP2B6, CYP2C8, CYP2C9, CYP2C19, CYP2D6, CYP3A4, CYP4A11, CES1, FMO3, and XDH), phase II metabolizing enzymes (e.g., GSTP1, NAT1, NAT2, TPMT, and members of the UGT1A family), and drug transporters (e.g., ABCB4, ABCC2, ABCC11, SLC10A2, SLC22A2, SLC22A12, SLC28A3, SLC2A9, SLC47A1, SLC47A2, and SLCO1B1). The panel also includes nuclear receptors regulating drug metabolism (NR1I2 and NR1I3), and genes involved in endogenous substrate metabolism or pharmacodynamic pathways, including CYP11B1, CYP17A1, CYP21A2, CYP24A1, DPYS, HNMT, and PNMT.

The second dataset, *D*_CYP_, is focused on a single ADME gene, CYP2C9, one of the most clinically relevant pharmacogenes.^18^ CYP2C9 encodes a phase I drug-metabolizing enzyme that plays a key role in the biotransformation of numerous clinically important drugs, including warfarin, phenytoin, nonsteroidal anti-inflammatory drugs, and sulfonylureas. Genetic variations in CYP2C9 give rise to normal, intermediate, or poor metabolizer phenotypes, which can substantially influence drug efficacy and toxicity. As a result, CYP2C9 genotype-guided prescribing recommendations have been incorporated into multiple guidelines for several drug classes.^4,19^ Two deep mutational scanning (DMS) datasets for CYP2C9, generated by,^20^ were collected from MaveDB.^21^ These datasets quantify the effects of all possible single amino acid variants on protein stability and enzymatic activity. To derive a single functional score for each variant, which we call DMS_fitness_, we combined the normalized stability and activity measurements, each ranging from 0 to 1, by multiplication, assuming that the overall functional impact is determined by the product of these two properties. Under this assumption, impairment of either protein stability or catalytic activity is sufficient to compromise the enzyme’s ability to metabolize its substrates. In this way, we generated the dataset D_CYP_, which contains functional characterization data for 8,091 non-synonymous variants of CYP2C9.

The third dataset, denoted as D_CPIC_, consists of 33 clinically annotated CYP2C9 star alleles with established functional assignments, as reported in recommendations issued by the Clinical Pharmacogenetics Implementation Consortium (CPIC); alleles with unknown or uncertain functions were excluded. The annotations were extracted from the supplementary material of.^20^

### 2.2. Benchmarked variant effect predictors

We evaluated several classes of computational predictors in our benchmark. First, we considered all 18 variant effect predictors previously benchmarked on dataset D_CL_ in.^17^ Based on their reported performance, we selected the eight best-performing predictors for further comparison: CADD,^22^ VEST3,^23^ MutationAssessor,^24^ PolyPhen-2,^8^ PROVEAN,^9^ SIFT,^10^ DANN^25^ and FATHMM-MKL.^26^ These methods represent widely used computational approaches for predicting the functional consequences of missense variants. We compared them with our in-house predictors FITMuSiC,^27^ RSALOR^15,28^ and StructureDCA,^16^ which assess variant effects on protein fitness by integrating evolutionary and structural information in different ways. Finally, we benchmarked several deep learning-based methods that have recently been proposed state of the art approaches for variant effect prediction; however, whether they truly represent the current state of the art remains debatable. In particular, we benchmarked AlphaMissense^13^ and four general pLLM models: ESM-2,^14^ SaProt,^29^ VenusREM,^30^ and ProSST.^31^

## 3. Results and Discussion

### 3.1. Assessing Method Performance on Experimental Intrinsic Clearance

We started by evaluating 16 variant effect predictors on the *D*_CL_ intrinsic clearance dataset, as described in Methods. Several observations emerge from the benchmark results presented in Table 1. First, the field has made substantial progress in recent years. The best-performing methods in the benchmark, namely StructureDCA, AlphaMissense, SaProt, and RSALOR, are all recent approaches. They achieved Spearman’s rank correlation coefficients *ρ* of approximately 0.65 between predicted scores and experimentally measured intrinsic clearance and consistently outperformed classical variant-effect predictors, including PROVEAN, PolyPhen-2, and SIFT, by more than 0.1. This substantial improvement demonstrates the advancement of the field in capturing the functional consequences of pharmacogenomic variants.

**Table 1.**
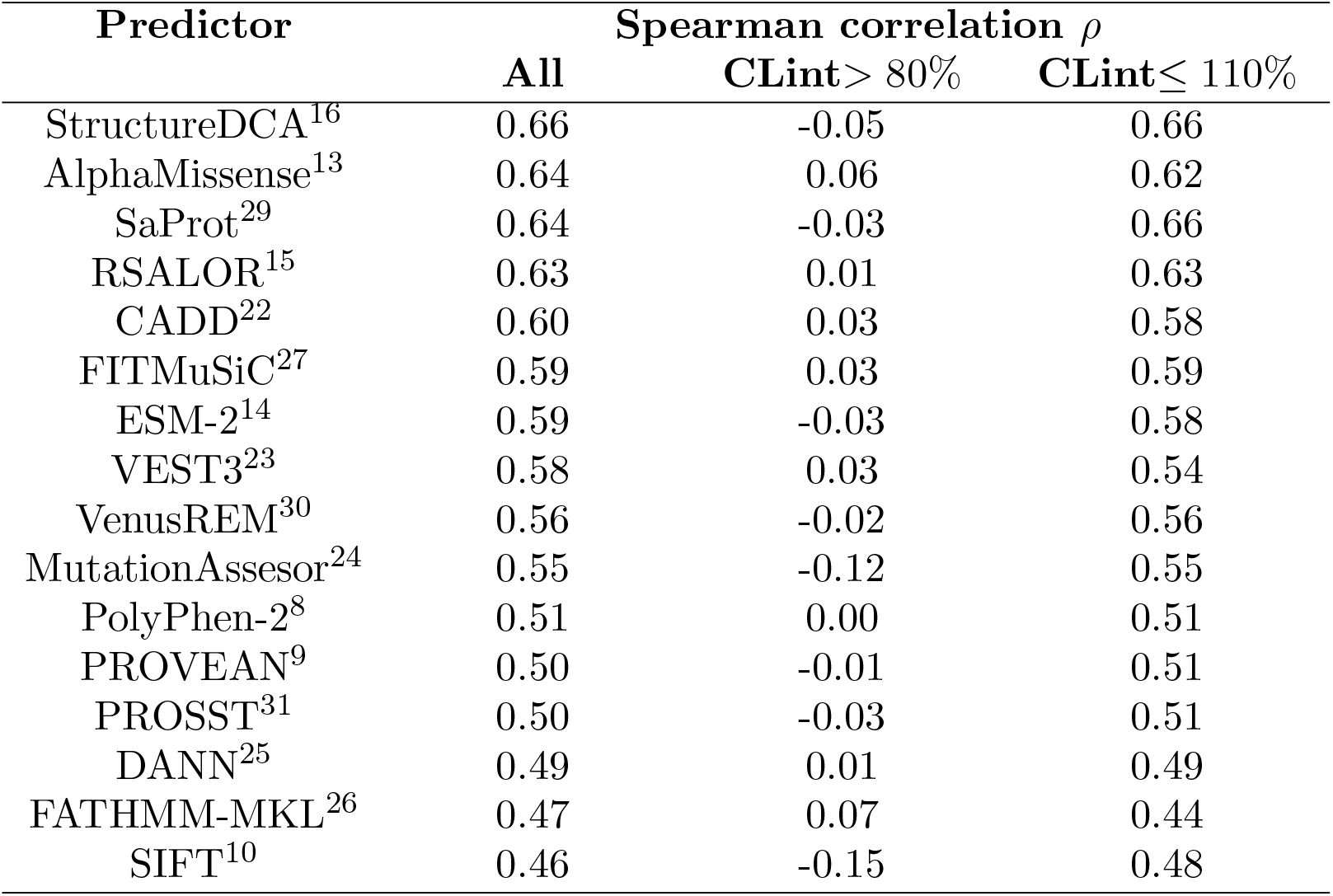
Benchmark performance on the *D*_CL_ dataset in terms of Spearman’s rank correlation coefficient (*ρ*) between predictor scores and experimentally measured intrinsic clearance (CLint), expressed as a percentage of the variant relative to the wild-type. Column 2 considers all variants; column 3 considers the subset of variants with CLint *>* 80%, comprising neutral and fast-metabolizing variants; and column 4 considers the subset with CLint ≤110%, comprising neutral variants and variants with impaired drug clearance.

| Predictor | Spearman correlation $\rho$ | | |
| --- | --- | --- | --- |
| | All | CLint $> 80\%$ | CLint $\leq 110\%$ |
| StructureDCA<sup>16</sup> | 0.66 | -0.05 | 0.66 |
| AlphaMissense<sup>13</sup> | 0.64 | 0.06 | 0.62 |
| SaProt<sup>29</sup> | 0.64 | -0.03 | 0.66 |
| RSALOR<sup>15</sup> | 0.63 | 0.01 | 0.63 |
| CADD<sup>22</sup> | 0.60 | 0.03 | 0.58 |
| FITMuSiC<sup>27</sup> | 0.59 | 0.03 | 0.59 |
| ESM-2<sup>14</sup> | 0.59 | -0.03 | 0.58 |
| VEST3<sup>23</sup> | 0.58 | 0.03 | 0.54 |
| VenusREM<sup>30</sup> | 0.56 | -0.02 | 0.56 |
| MutationAssesor<sup>24</sup> | 0.55 | -0.12 | 0.55 |
| PolyPhen-2<sup>8</sup> | 0.51 | 0.00 | 0.51 |
| PROVEAN<sup>9</sup> | 0.50 | -0.01 | 0.51 |
| PROSST<sup>31</sup> | 0.50 | -0.03 | 0.51 |
| DANN<sup>25</sup> | 0.49 | 0.01 | 0.49 |
| FATHMM-MKL<sup>26</sup> | 0.47 | 0.07 | 0.44 |
| SIFT<sup>10</sup> | 0.46 | -0.15 | 0.48 |

The second insight gained from these results is that not all recent methods perform equally well and that the best-performing methods are not necessarily the most complex. A striking example is our in-house tool RSALOR, a very simple model that combines the residue solvent accessibility (RSA) of the mutated residue and the log-ratio of site-independent sequence conservation. As shown in Table 1, it outperforms several considerably more complex models comprising millions of parameters and requiring extensive training. In particular, methods that rank among the top performers on the ProteinGym benchmark, such as the pLLM model VenusREM that incorporates structural and evolutionary information, perform worse on our pharmacogenomic benchmark. These findings suggest that increasing model complexity does not necessarily translate into better predictive performance.

The last insight that emerges from this benchmark comes from the last two columns of Table 1, which compare the ability of the predictors to distinguish between neutral variants and variants associated with increased metabolic activity (column 3) and between neutral and loss-of-function variants that impair drug clearance (column 4). This highlights one of the major challenges in the field of computational variant effect prediction. Indeed, predicting variants that increase drug clearance relative to the wild-type remains extremely difficult for all methods, with Spearman’s rank correlation coefficients close to zero on the subset containing neutral and increased-clearance variants. In contrast, the predictors perform considerably better at identifying loss-of-function variants that impair drug clearance. This is likely because evolutionary information is much more informative for detecting deleterious substitutions, as non-functional variants are generally eliminated by natural selection and are therefore rarely observed during evolution, resulting in strong conservation signals.

### 3.2. Assessing Method Performance on Experimental CYP2C9 Fitness

Next, we evaluated variant effect on the D_CYP_ dataset, comprising DMS data on CYP2C9 variants. The CYP2C9 gene encodes cytochrome P450 2C9, one of the major phase I drug-metabolizing enzymes involved in the biotransformation of numerous clinically important drugs.^32,33^ The enzyme adopts the canonical cytochrome P450 fold, formed by an N-terminal transmembrane helix that anchors the protein to the endoplasmic reticulum membrane, and a catalytic domain comprising a large predominantly *α*-helical subdomain and a small *β*-sheet-rich subdomain.

In a recent investigation,^20^ a comprehensive DMS map of all CYP2C9 variants was generated, measuring both stability and enzymatic activity. We combined these two measurements into a single fitness score called DMS_fitness_ by multiplying the two experimental scores (see Methods). Here, we benchmarked the four best-performing methods identified in the previous subsection (Table 1) to investigate their respective strengths and weaknesses in greater detail.

Our coevolution-based method StructureDCA, which integrates wild-type structural information, achieves the best overall performance among the four tested approaches, reaching a correlation coefficient *ρ* of 0.66. Notably, our simple site-independent evolution-based model, RSALOR, also performs remarkably well, achieving better performance than AlphaMissense, a deep learning method that reuses AlphaFold^34^ representations and is often considered as a state-of-the-art approach. Moreover, the results presented in Table 2 are consistent with those obtained on the D_CL_ dataset (Table 1): the correlation is substantially higher when distinguishing neutral from loss-of-function variants than when distinguishing neutral from gain-of-function variants. Nevertheless, unlike our observations for the D_CL_ dataset, the four predictors achieve a lower but still statistically significant correlation for the neutral versus gain-of-function subset, indicating that they are able to capture, to some extent, the molecular determinants underlying increased enzymatic activity.

**Table 2.** Benchmark performance on the *D*_CYP_ dataset in terms of Spearman’s rank correlation coefficient *ρ* between the predictors’ scores and the measured DMS_fitness_ scores defined in Methods. Column 2 considers all variants; column 3 considers the subset of variants with DMS_fitness_ *>* 0.8, comprising neutral and hyper-active variants ; and column 4 considers the subset with DMS_fitness_ ≤ 1.1, comprising neutral and inactive variants.

| Predictor | Spearman correlation $\rho$ | | |
| --- | --- | --- | --- |
| | All | $\text{DMS}_{\text{fitness}} > 0.8$ | $\text{DMS}_{\text{fitness}} \leq 1.1$ |
| StructureDCA<sup>16</sup> | 0.66 | 0.24 | 0.64 |
| SaProt<sup>29</sup> | 0.64 | 0.21 | 0.62 |
| RSALOR<sup>15</sup> | 0.63 | 0.21 | 0.62 |
| AlphaMissense<sup>13</sup> | 0.61 | 0.13 | 0.61 |

To better understand the strengths and limitations of StructureDCA, the best-performing predictor in both benchmarks, we performed a more detailed analysis across different structural regions and variant classes. First, we computed the mean StructureDCA and DMS_fitness_ scores across all substitutions at each residue position. As shown in Fig. 1, the correlation is even stronger at the residue level, with Spearman’s rank correlation coefficient reaching *ρ* = 0.73. We also analyzed predictive performance across the different structural and functional regions of the CYP2C9 enzyme. The transmembrane helix that anchors the enzyme to the membrane is connected to the catalytic domain by a 15-residue linker. The catalytic domain comprises several key elements, including the six substrate-recognition sites (SRS1–SRS6), which line the active-site cavity and determine substrate binding, orientation, and catalytic specificity; the highly conserved heme-binding motif and the adjacent meander region, an extended loop between helices K and L that is essential for maintaining the structural integrity of the catalytic center; the JK region, comprising helices J and K and the intervening JK loop, which interacts with the meander region to stabilize the catalytic core; and several flexible loop regions, notably the GH loop between helices G and H and the CD and EF loops (CD/EF), which connect helices C and D and helices E and F, respectively.

**Fig. 1.**
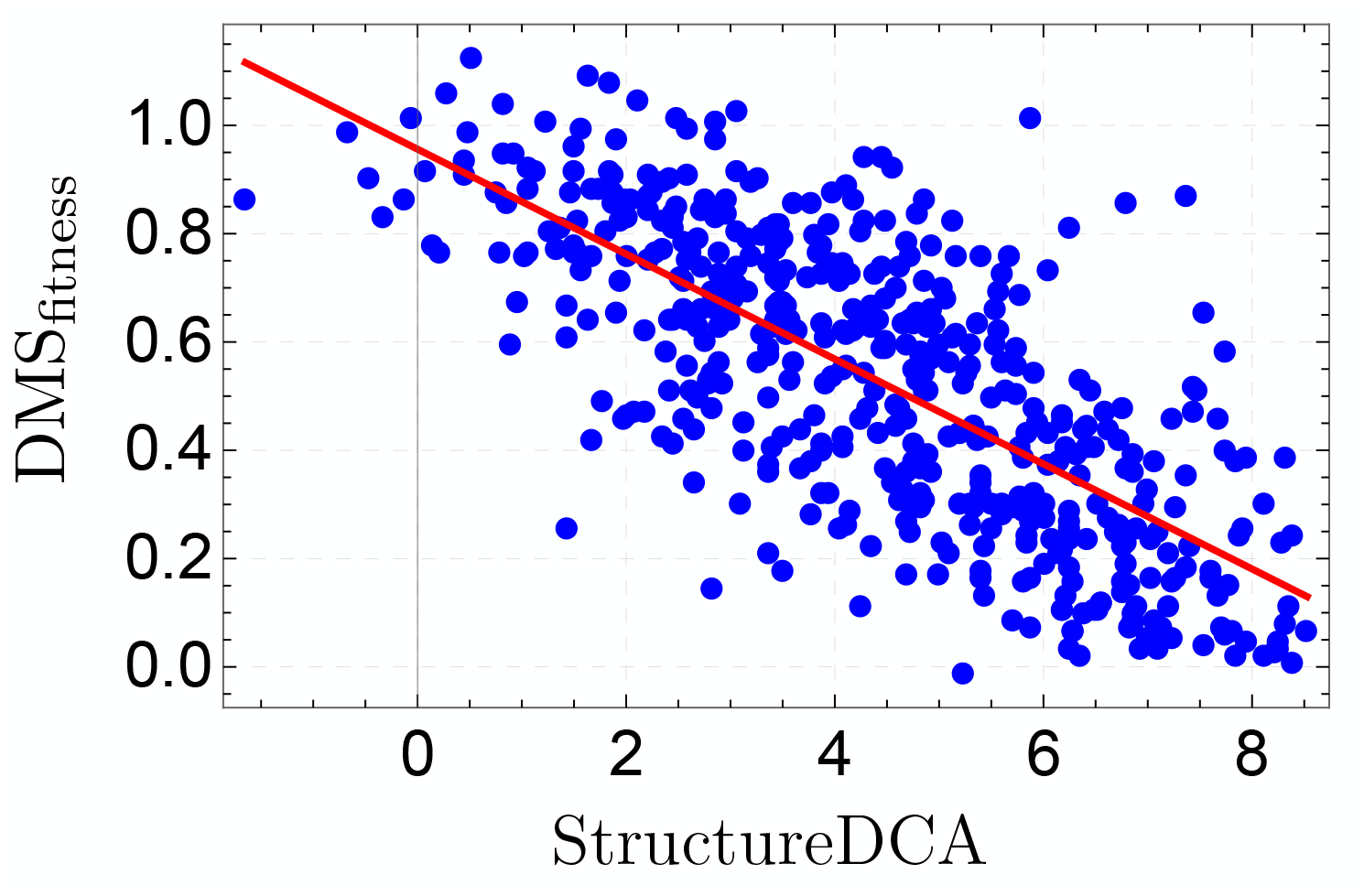
The mean per-residue DMS_fitness_ scores as a function of the mean per-residue StructureDCA scores for all CYP2C9 variants in the D_CYP_ dataset.

As shown in Table 3, the highest correlations are generally observed in highly conserved regions, such as the heme-binding region and the JK region. However, some regions with relatively low conservation, such as the linker region, are also predicted remarkably well. In contrast, the performance is substantially lower for the substrate-recognition sites, which are primarily responsible for substrate recognition and specificity rather than structural stability, indicating that mutations affecting them remain considerably more challenging to interpret. Although variants within the substrate-recognition sites may have only a limited impact on protein fitness and are therefore subject to weaker evolutionary constraints, they can have profound functional and clinical consequences by selectively altering the metabolism of specific drugs. For example, the I359L substitution in SRS5 reduces the hydroxylation of tolbutamide and phenytoin while leaving the diclofenac metabolism largely unaffected.^35,36^ Similarly, the F114L substitution in SRS1 impairs warfarin hydroxylation and reduces inhibition by sul-

**Table 3.**
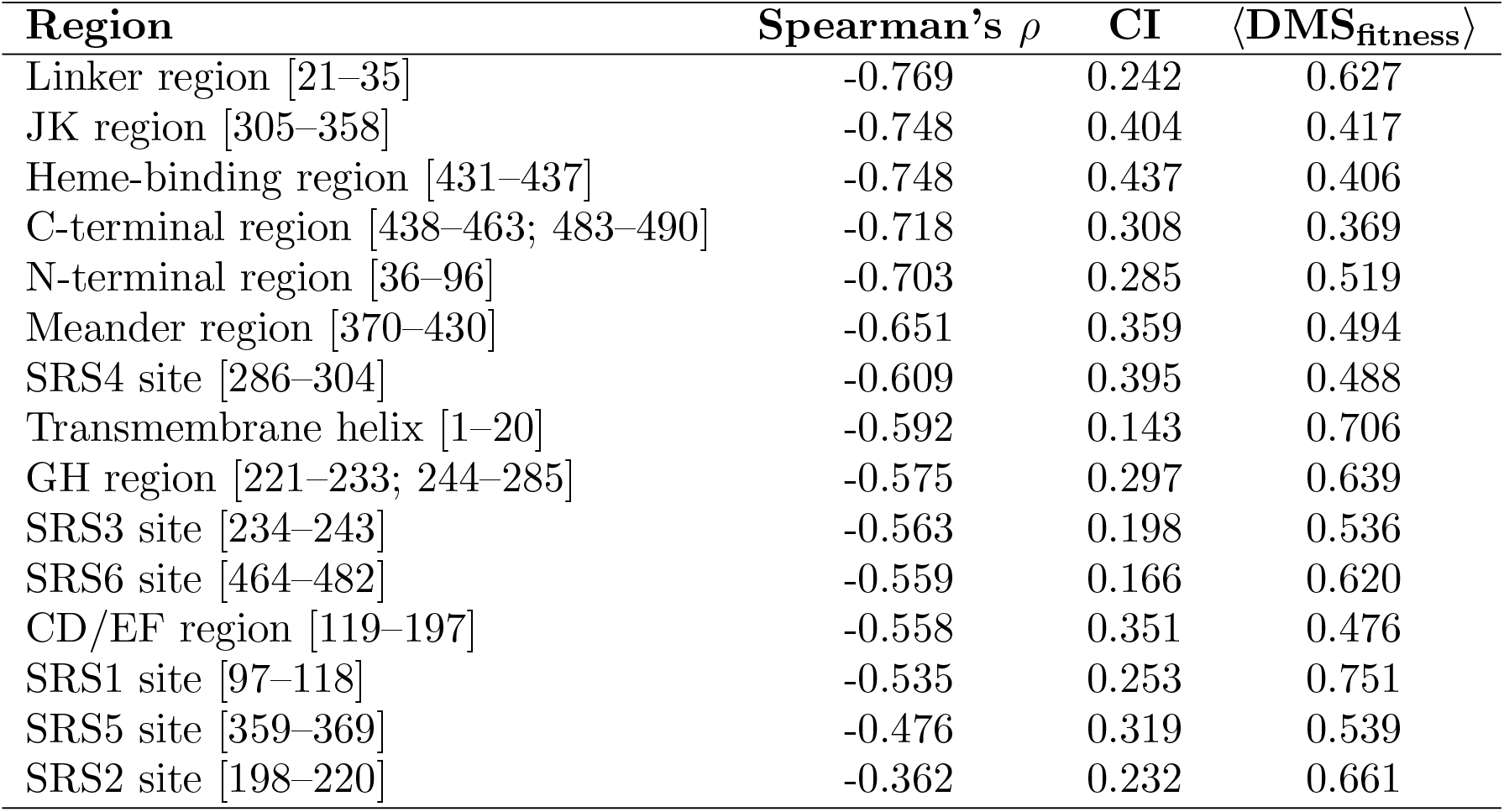
Spearman’s rank correlation coefficient (*ρ*) between the mean StructureDCA score and the mean experimental DMS_fitness_ score across the different structural regions of CYP2C9, for variants in the *D*_CYP_ dataset; the residue boundaries are provided in brackets. The third column reports the mean conservation index (CI) for each region, and the fourth column the mean DMS score (⟨DMS_fitness_⟩). Regions are ranked from highest to lowest absolute correlation |*ρ*|.

### 3.3. Assessing Method Performance on CYP2C9 Clinical Annotations

Finally, we evaluated the performance of the StructureDCA predictor on clinically annotated CYP2C9 variants to assess their ability to capture clinically relevant functional effects. More precisely, we compared the predictions of StructureDCA as well as the experimental DMS measurements with the functional annotations assigned by the Clinical Pharmacogenetics Implementation Consortium (CPIC), which provides evidence-based guidelines linking CYP2C9 star alleles to their expected functional activity and corresponding therapeutic recommendations;^19,38,39^ these variants were compiled in the *D*_CPIC_ dataset.

The clinical annotations classify variants into three categories: variants with wild-type function (denoted by =), variants with decreased function (↓), and variants with no function (∅). For comparison, we defined corresponding functional categories for the DMS_fitness_ and StructureDCA scores. Variants with DMS_fitness_ scores below 0.3 and above 0.8 were classified as having abolished function and wild-type function, respectively; variants with intermediate scores (0.3–0.8) were assigned to the decreased-function category. To define the corresponding thresholds for StructureDCA, we used the linear relationship between DMS_fitness_ and

StructureDCA scores shown in Fig. 1. Specifically, the StructureDCA scores corresponding to DMS_fitness_ values of 0.3 and 0.8 on the fitted regression line were used as the StructureDCA thresholds. As shown in Table 4, both the experimental and computational approaches exhibit reasonable overall performance. However, neither achieves high accuracy, as both misclassify several clinically annotated variants. Interestingly, StructureDCA predictions are more consistent with the clinical phenotype than the DMS scores, with an accuracy of 66.7% compared with 50% for DMS_fitness_ scores.

**Table 4.** Comparison between CPIC functional annotations, StructureDCA prediction scores and experimental DMS_fitness_ scores for CYP2C9 alleles collected in the *D*_CPIC_ dataset. The symbols =, ↓and ∅ denote variants with wild-type, decreased, and abolished function, respectively.

| Allele | Mutation | Function | StructureDCA | DMS<sub>fitness</sub> |
| --- | --- | --- | --- | --- |
| *2 | R144C | ↓ | ↓ | ↓ |
| *3 | I359L | ↓ | ↓ | ↓ |
| *4 | I359T | ↓ | ↓ | = |
| *5 | D360E | ↓ | ↓ | - |
| *8 | R150H | ↓ | = | ↓ |
| *9 | H251R | = | ↓ | = |
| *11 | R335W | ↓ | ↓ | ∅ |
| *12 | P489S | ↓ | ∅ | ∅ |
| *13 | L90P | ↓ | ↓ | ∅ |
| *14 | R125H | ↓ | ↓ | ↓ |
| *16 | T299A | ↓ | ↓ | ↓ |
| *23 | V76M | ↓ | ↓ | ∅ |
| *24 | E354K | ∅ | ∅ | ∅ |
| *26 | T130R | ↓ | ↓ | = |
| *28 | Q214L | ↓ | ↓ | ↓ |
| *29 | P279T | ↓ | = | - |
| *30 | A477T | ↓ | = | ↓ |
| *31 | I327T | ↓ | ↓ | ∅ |
| *33 | R132Q | ↓ | ↓ | - |
| *37 | D49G | ↓ | ↓ | = |
| *38 | G96A | ↓ | ↓ | = |
| *39 | G98V | ∅ | ↓ | ↓ |
| *42 | R124Q | ∅ | ↓ | ↓ |
| *43 | R124W | ∅ | ∅ | ∅ |
| *44 | T130M | ↓ | ↓ | ↓ |
| *45 | R132W | ∅ | ↓ | ∅ |
| *46 | A149T | ↓ | ↓ | ∅ |
| *50 | P227S | ↓ | ↓ | ↓ |
| *52 | T299R | ∅ | ↓ | ∅ |
| *55 | L361I | ↓ | = | ↓ |
| *58 | P337T | ↓ | ↓ | ∅ |
| *59 | I434F | ↓ | ↓ | = |
| *60 | L467P | ↓ | = | ∅ |
| Accuracy |  |  | 22/33<br>66.7% | 15/30<br>50.0% |

Let us examine some variants to understand why and where the method fails. One example is the CYP2C9*8 allele carrying the R150H mutation, which StructureDCA predicts to have no functional effect, whereas it is correctly classified by the DMS score. This variant is located in the CD/EF helical region and is exposed on the protein surface. StructureDCA likely predicts it as neutral because the position is poorly conserved and solvent-exposed. In contrast, the DMS_fitness_ score is approximately 0.5 on a scale from 0 to 1, indicating a mild detrimental effect on both activity and stability. R150H defines the CYP2C9*8 allele, which is found predominantly in individuals of African ancestry. It has been associated with lower warfarin dose requirements in African–American patients to achieve an optimal anticoagulation response.^40^

In contrast, the CYP2C9*26 allele carrying the T130R mutation shows reduced function, with multiple in vitro studies demonstrating decreased clearance relative to the wild type across different cell systems and substrates, including warfarin, tolbutamide, diclofenac, losartan, and glimepiride.^41,42^ This effect is correctly captured by StructureDCA, whereas the DMS score indicates an almost neutral impact and therefore fails to reproduce the clinically observed phenotype.

Taken together, these results highlight an important point that is not always fully taken into consideration: the direct translation of in vitro measurements into clinical phenotypes is not straightforward, and high-performing computational methods may complement experimental data, better reflect the observed phenotype in some cases, and help variant interpretation.^12^

## 4. Conclusion

Despite recent advances in computational approaches, the interpretation of pharmacogenomic variants remains an open challenge and a critical step toward the realization of personalized medicine. Even the best-performing methods achieve only moderate correlations with experimentally measured intrinsic clearance, indicating that a substantial fraction of phenotypic variability remains unexplained. This limitation arises because variant effects depend on multiple factors beyond protein stability and evolutionary conservation, including substrate specificity, catalytic mechanisms, conformational dynamics, and allosteric effects, all of which are difficult to predict.

A major limitation of the predictors evaluated in this study is their ability to identify gain-of-function variants that increase pharmacogenomic clearance through enhanced enzymatic activity. All methods showed limited, or virtually no, capacity to distinguish these variants from neutral variants and consequently failed to identify fast-metabolizer phenotypes. These findings highlight substantial opportunities for further improvement.

Another key message emerging from our analysis is that recent advances in state-of-the-art variant interpretation methods do not always translate into substantial gains in predictive performance, despite an overall improvement compared with classical approaches. For example, the simple model RSALOR,^15^ which relies primarily on evolutionary conservation and solvent accessibility, achieves comparable or superior predictive performance than AlphaMissense and recent pLLMs. The best-performing model in our analysis, StructureDCA, combines physical principles with evolutionary constraints within an interpretable framework, suggesting that biophysically informed integration may represent a promising strategy compared with black-box artificial intelligence-based models trained on huge datasets.

Finally, by comparing DMS values with StructureDCA scores for predicting functional phenotypes associated with the clinically important CYP2C9 enzyme, we further demonstrated that computational methods, despite their limitations, can play a fundamental role in clinical variant-effect interpretation. These approaches may help resolve cases in which in vitro experiments are inconclusive or fail to fully capture the observed clinical phenotype, as previously discussed in the field.^12^

## Notes

### Competing Interest Statement

The authors have declared no competing interest.

### Funding

- FNRS Belgique

